# Hypercaloric diet triggers transient molecular rearrangements of astrocytes selectively in the arcuate nucleus

**DOI:** 10.1101/2022.03.30.486358

**Authors:** Luiza Maria Lutomska, Viktorian Miok, Natalie Krahmer, Ismael González García, Tim Gruber, Ophélia Le Thuc, Cahuê De Bernardis Murat, Beata Legutko, Michael Sterr, Gesine Saher, Heiko Lickert, Timo D. Müller, Siegfried Ussar, Matthias H. Tschöp, Dominik Lutter, Cristina García-Cáceres

## Abstract

Hypothalamic astrocytes are particularly affected by energy-dense food consumption. How the anatomical location of these glial cells and their spatial molecular distribution in the arcuate nucleus of the hypothalamus (ARC) determine the cellular response to a high caloric diet remains unclear. In this study, we investigated their distinctive molecular responses following the exposure to a high-fat high-sugar (HFHS) diet, specifically in the ARC. Using RNA sequencing and proteomics, we showed that astrocytes have a distinct transcriptomic and proteomic profile dependent on their anatomical location, with a major proteomic reprogramming in hypothalamic astrocytes. By ARC single-cell sequencing, we observed that a HFHS diet dictates time- and cell-specific transcriptomic responses, revealing that astrocytes have the most distinct regulatory pattern compared to other cell types. Lastly, we topographically and molecularly characterized astrocytes expressing glial fibrillary acidic protein and/or aldehyde dehydrogenase 1 family member L1 in the ARC, of which the abundance was significantly increased, as well as the alteration in their spatial and molecular profiles, with a HFHS diet. Together, our results provide a detailed multi-omics view on the spatial and temporal changes of astrocytes particularly in the ARC during different time points of adaptation to a high caloric diet.

## INTRODUCTION

Astrocytes have been historically classified based on their morphology, anatomical location, and marker gene expression (Barres, 2008, Sofroniew and Vinters, 2010). Indeed, canonical astrocyte markers do not always label the entire population of astrocytes and/or the same embedded circuitries (Kimelberg, 2004, Emsley and Macklis, 2006, Preston et al., 2019). Recent studies based on bulk and single-cell (sc) RNA sequencing (Seq) analyses have revealed that astrocytes exhibit a high molecular diversity within the same, as well as across different brain regions (Ben Haim and Rowitch, 2017, Morel et al., 2017, Boisvert et al., 2018). Moreover, astrocytes show a selective region-dependent vulnerability to specific diseases (Barres, 2008, Seifert et al., 2006). Regarding metabolic control, multiple studies have indicated region-specific roles of astrocytes in the regulation of neuronal circuitries, as indicated by the fact that selective perturbations in astrocytes trigger a wide array of alterations in energy balance and glucose homeostasis (Garcia-Caceres et al., 2016, Gao et al., 2017, Stein et al., 2020). In response to hypercaloric diet, astrocytes located in the arcuate nucleus of the hypothalamus (ARC) undergo morphological modifications characterized by a hypertrophic phenotype and an increase in glial fibrillary acidic protein (GFAP) immunoreactivity (Horvath et al., 2010, Thaler et al., 2012, Sofroniew, 2014, Escartin et al., 2021). This astrogliosis occurs in the ARC of both obese humans and mice (Thaler et al., 2012), affecting the cytoarchitecture and synapses of melanocortin circuits, and altering their physical interactions with neurons and blood vessels in mice (Horvath et al., 2010). Notably, diet-induced increased gliosis in the ARC was observed prior to body weight gain and/or peripheral signs of inflammation in murine models (Thaler et al., 2012, Buckman et al., 2015). However, it remains to be elucidated whether hypothalamic astrocytes, and particularly ARC astrocytes, exhibit a distinct specialization based on their distribution and marker expression, which might confer the selective sensitivity to respond to metabolic diseases.

In this study, we investigated the transcriptional and proteomic response of astrocytes to a long-term hypercaloric diet exposure. We found that astrocytic response depends on anatomical location, which was reflected by changes in the hypothalamic proteome that exceeded those in the hippocampus and cortex. Moreover, as indicated by our scRNA-Seq analysis, a short exposure to a high-fat high-sugar (HFHS) diet (5 days) was sufficient to induce astrocyte-specific transcriptional changes in the ARC. In contrast to other ARC cell types, these transcriptional changes were widely reduced after 15d exposure to a HFHS diet. Furthermore, we identified that astrocytes expressing aldehyde dehydrogenase 1 family member L1 (Aldh1L1) and/or GFAP increase in number depending on spatial positions, and change their transcriptional profile in the ARC in response to a HFHS diet feeding. Overall, our results highlight specific gene and protein profiles of hypothalamic astrocytes in response to a HFHS diet, as well as show that ARC astrocytes are characterized by distinctive temporal, transcriptional and molecular rearrangements in response to a hypercaloric diet.

## RESULTS

### The anatomical location of astrocytes strongly influences their sensitivity to a HFHS diet

We examined the gene expression of purified astrocytes collected from the hypothalamus, hippocampus, and cortex of wildtype adult male mice exposed to a standard chow (SC) diet or a HFHS diet for 4 months. Magnetic-activated cell sorting (MACS) was employed to isolate astrocytes from the dissected brain areas (Fig. 1A) by using astrocyte cell surface antigen 2 (ACSA2) as marker (Batiuk et al., 2017, Kantzer et al., 2017). The samples were subjected to bulk RNA-Seq and proteomics analyses, including the unlabeled fraction (ACSA2^-^) as a negative control for proteomics (Fig. 1A-B). We found that ACSA2^+^ cells showed a high abundancy of canonical astrocyte markers, such as *Aldh1l1*, Aquaporin 4 (*Aqp4*), *Gfap*, solute carrier family 1 member 2 (*Slc1a2*), and solute carrier family 1 member 3 (*Slc1a3*) (Fig. 1B). Consistent with previous observations (Kantzer et al., 2017), some markers for oligodendrocytes (*Mag, Mog*) and mural cells (*Des, Mustn1, Pdgfrb*) were detected in ACSA2^+^ cells in transcriptomics and proteomics analyses, respectively. In contrast, ACSA2^-^ cells showed a high enrichment of neuronal (*Snap25, Syp, Syt1, Tubb3*) and microglia (*Aif1, Itgam*) specific markers (Fig. 1B). Next, we performed a principal component analysis (PCA) of both transcriptome and proteome datasets derived from ACSA2^+^ fractions, which showed that the expression profile of astrocytes depends on their anatomical location rather than on the effect of the diet (Fig. 1C). We also identified the differentially expressed genes (DEGs) in specific region-derived astrocytes from mice fed with a SC or a HFHS diet (Fig. S1A-B). Notably, we found only 12 overlapping DEGs in response to a HFHS diet in the three brain regions analyzed (Fig. S1A and Table S1). Conversely, we identified 2514, 868, and 563 non-overlapping DEGs in the cortex, hypothalamus, and hippocampus, respectively (Fig. S1A and Table S1). To further identify molecular pathways potentially modulated by a hypercaloric diet, we performed GO enrichment analysis on the non-overlapping DEGs from the hierarchical clustering (Fig. S1C and Table S1). Our data show that consumption of a calorie-rich diet mainly down-regulated genes encoding proteins involved in metabolically relevant pathways in cortical astrocytes, such as “*insulin receptor signaling pathway*,” “*response to insulin*,” and “*TOR signaling*,” and pathways related to inter-cellular organization (“*cell-matrix adhesion*”), and microvasculature (“*regulation of angiogenesis*”) in hippocampal astrocytes. In hypothalamic astrocytes, long-term exposure to a HFHS diet induced an up-regulation of DEGs enriched in relevant pathways related to glucose metabolism (“*pyruvate metabolic process*”), lipid metabolism (“*fatty acid metabolic process*” and “*fatty acid oxidation*”), and metabolic stress (*“response to reactive oxygen species”)*, and a down-regulation of pathways involved in neuronal and synaptic regulation (“*synaptic transmission, GABAergic*,” “*AMPA glutamate receptor clustering*,” and “*regulation of synapse structure or activity*”) (Fig. S1C and Table S1). Given that alterations in protein abundance can influence cell function without entailing transcriptional changes (Liu et al., 2016, Buccitelli and Selbach, 2020), we analyzed the astrocytic proteome. Using a similar approach as detailed above, we identified differentially expressed proteins (DEPs) and performed the GO enrichment (Fig. 1D-F). We identified 447 unique DEPs in the hypothalamus, compared to 92 in the cortex, and 124 in the hippocampus, without observing common DEPs between these three brain regions (Fig. 1D). The GO enrichment analysis on the non-overlapping DEPs (Fig. 1F and Table S2) revealed that cortical astrocytes showed mainly up-regulated glucose metabolism- (“*cellular response to glucose stimulus*”) and metabolic stress- (“*response to reactive oxygen species*”) associated pathways, and down-regulated proliferation- (“*positive regulation of G1/S transition of mitotic cell cycle*”) associated pathway. In hippocampal-isolated astrocytes, the DEPs were mainly down-regulated in pathways related to glucose metabolism (“*cellular carbohydrate metabolic process*”), proliferation (“*smoothened signaling pathway*”), kinetics of vesicle release (“*calcium ion transport into cytosol*”), and neuronal and synaptic regulation (“*axon guidance*”). Interestingly, the proteome of hypothalamic astrocytes was remarkably affected by a HFHS diet with DEPs significantly up-regulated in pathways related to lipid and hormonal signaling regulation (“*fatty acids catabolic process*,” “*insulin receptor signaling pathway*,” “*leptin mediated signaling pathway*,” and “*response to leptin*”). Moreover, hypothalamic astrocyte-specific DEPs were enriched in other crucial pathways, with an overall down-regulation in glucose metabolism- (“*glycogen metabolic process*” and “*glucose transmembrane transport*”), kinetics of vesicle release- (“*calcium-mediated signaling*”), and neuronal and synaptic regulation- (“*regulation of neurotransmitter levels*,” “*glutamate receptor signaling pathway*,” and “*regulation of neuronal synaptic plasticity*”) related pathways (Fig. 1F). Comparing transcriptomic and proteomic expression, we did not observe a significant correlation between transcripts and related proteins on molecular level. However, we could confirm several previously identified pathways enriched for significantly co-upregulated (“*TOR signaling*,” “*glycolytic process*,” “*cellular response to glucose stimulus*,” and “*response to reactive oxygen species*”) or co-down-regulated (“*regulation of mitotic cell cycle*”) transcript-protein pairs in cortical astrocytes. Unlike hippocampus, in the astrocytes of which no common up- or down-regulated pathways were detected, hypothalamic astrocytes showed common up-regulated pathways (“*fatty acid metabolic process*,” “*fatty acid catabolic process*,” and “*fatty acid oxidation*”) (Fig. S1D and Table S3). The lack of correlation between proteomics and transcriptomics is a frequently reported observation and subject of ongoing debate (de Sousa Abreu et al., 2009, Payne, 2015). Overall, these findings indicate that long-term exposure to hypercaloric diet induces molecular changes in astrocytes in a region-dependent manner. In particular, the major proteomic switch was observed in hypothalamic astrocytes, specifically in pathways related to nutritional and hormonal signaling regulation.

**Figure 1.**
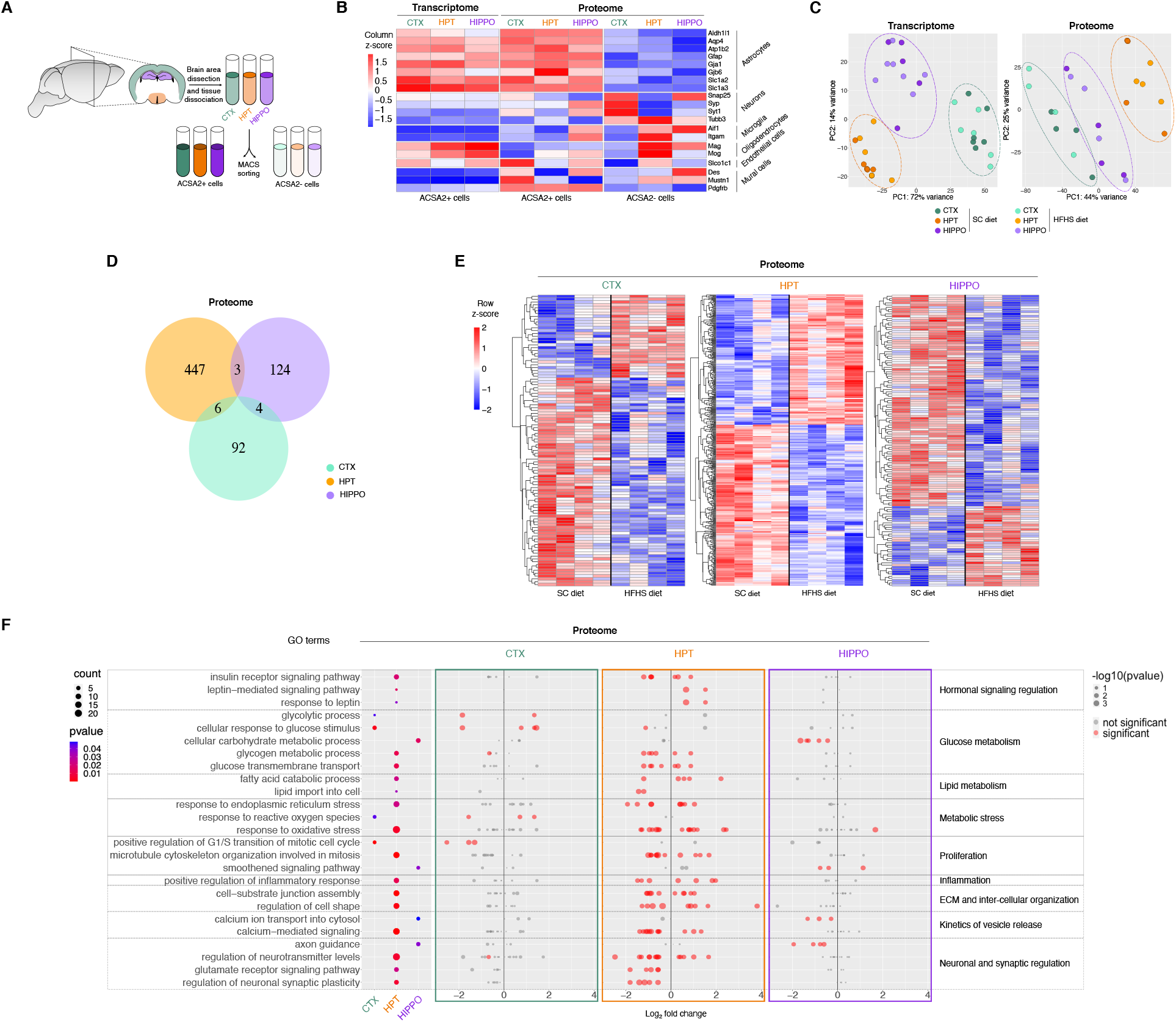
MACS-isolated astrocytes from distinct brain regions reveal transcriptional heterogeneity in response to a HFHS diet. **(A)** Schematic describing the isolation of ACSA2^+^ and ACSA2^-^ cells from cortex (CTX, green), hypothalamus (HPT, orange) and hippocampus (HIPPO, purple) of mice. Brain areas were dissected and dissociated into single-cell suspensions and subjected to magnetic-activated cell sorting (MACS) to separate ACSA2^+^ cells from ACSA2^-^ cells. **(B)** Mean expression of diverse CNS cell type markers in different brain areas (cortex, hypothalamus, and hippocampus) in ACSA2^+^ (transcriptome and proteome), and ACSA2^-^ (proteome) cells, represented as column z-scores. The cells were isolated from mice fed with a SC diet or a HFHS diet for 4 months. Per each sample, 2 or 4 mice were combined for bulk RNA-Seq and proteomics analyses respectively. A number of 5-7 replicates (transcriptomics) and 4 replicates (proteomics) per each group were used. The datasets derived from SC diet and HFHS diet fed mice were combined for the analysis. **(C)** Principal component analysis of transcriptome and proteome derived from cortical, hypothalamic and hippocampal ACSA2^+^ astrocytes isolated from mice fed with a SC diet or 4 months of a HFHS diet. Each dot represents one sample. Each sample is formed by the combination of 2 (transcriptomics) or 4 (proteomics) mice. The dashed lines surround the samples derived from the same brain region. **(D)** Venn diagram depicting the numbers of commonly and differentially regulated proteins with p < 0.05 in ACSA2^+^ cells, comparing SC and HFHS diet groups in each brain region. **(E)** Heatmap of normalized proteome intensity values obtained by unsupervised clustering of differentially expressed proteins (p < 0.05) for each brain region. The color code in the heatmap indicates row z-score normalized expression values. Samples derived from SC or HFHS diet-fed animals are separated by a black line. Each column represents one sample, and each sample is formed by ACSA2^+^ cells derived from 4 animals. **(F)** The dot plot illustrates manually selected GO terms of interest enriched for DEPs identified in brain regions comparing SC and HFHS diet groups, in proteomics (p < 0.05). The left panel shows the dot plot of the selected GO enriched biological process terms for each brain tissue. The dot size indicates the number of DEPs, while the color depicts the statistical significance of the enrichment. The dot plots in the three adjacent panels correspond to the three different brain regions analyzed, where each protein mapped to a specific pathway is represented by a single dot. The dot color refers to the presence (red) or absence (grey) of a statistically significant value, and the dot size indicates the level of significance. The x-axis represents log_2_ fold change. ACSA2: astrocyte cell surface antigen 2; CTX: cortex; ECM: extracellular matrix; GO: gene ontology; HFHS: high-fat high-sugar; HIPPO: hippocampus; HPT: hypothalamus; MACS: magnetic-activated cell sorting; SC: standard chow. P values for differential protein expression were analyzed by Student’s *t*-test.

### ARC astrocytes exhibit distinctive temporal HFHS diet-induced transcriptional changes

Given that the ARC orchestrates whole-body energy metabolism (Jais and Brüning, 2021, González-García and García-Cáceres, 2021), and the morphology of ARC astrocytes is particularly affected by a hypercaloric diet (Thaler et al., 2012), we assessed the effects of a HFHS diet feeding over time on ARC-derived cells, to investigate their diet- and time-dependent transcriptomic modifications. Based on previous studies from our group showing that 15 days (d) of a HFHS diet exposure are sufficient to increase the body weight of mice in comparison to a standard chow (SC) diet (Gruber et al., 2021), we selected two critical time points: 5 and 15d of exposure to a HFHS diet for the assessment. We then performed scRNA-Seq on the totality of cells obtained from ARC dissections (21143 cells, of which 19995 were analyzed after filtering, divided in SC diet = 6741 cells; 5d HFHS diet = 5886 cells; 15d HFHS diet = 7368 cells; a median of 1802 unique transcripts was detected per each cell). Leiden clustering of the scRNA-Seq data identified 14 distinct clusters, which were mapped into 9 different cellular identities based on the expression of selected cell type-specific canonical markers (Fig. 2A-C). Astrocytes showed the highest number of DEGs (the majority of which were up-regulated) after 5d exposure to a HFHS diet, which then reversed to normal levels after 15d, when compared to the SC diet-fed group (starting point: 0 DEGs) (Fig. 2D, 2E and Table S4). In contrast, the other cell types exhibited the greatest number of DEGs (the majority of which were down-regulated) at 15d HFHS diet (Fig. 2D and 2E). Surprisingly, according to the number of DEGs, microglia and mural cells were not particularly affected at the selected time points of exposure to a HFHS diet (Fig. 2D and 2E). Therefore, the time of exposure to a HFHS diet influences transcriptomic responses in the ARC in a cell type-specific manner, in which astrocytes exhibit a distinct pattern of change in comparison to neighboring cells. We next investigated whether a HFHS diet induces ARC specific changes in the kinetics of gene expression, by studying the transcriptome of ARC astrocytes at two different time points after diet switch. For this, we first performed uniform manifold approximation and projection (UMAP) on astrocytes, and subsequently Leiden clustering on the dietary-wise subdivided cell projections (Fig. 2F). We identified four sub-clusters of astrocytes in SC diet and 15d HFHS diet experimental groups, and three sub-clusters in 5d HFHS diet group (Fig. 2F). Of note, the sub-cluster *3* in the 15d HFHS diet group was comprised by only 15 cells and was not considered in the subsequent analyses. Interestingly, the three remaining sub-clusters of the 15d HFHS diet group appeared to be in great consensus with the three sub-clusters of the 5d HFHS group, thus we labelled these sub-clusters accordingly as *0, 1*, and *2* (Fig. 2F). Moreover, sub-cluster *a* was split in sub-clusters *0* and *1*, while the three sub-clusters *b, c*, and *d* were combined in sub-cluster *2*, under a HFHS diet (Fig. 2F). To study how transcriptional activity may explain this shift in the clustering, we performed RNA velocity analysis on the astrocyte sub-clusters (La Manno et al., 2018, Bergen et al., 2020). We observed that the transcriptional activity of astrocytes derived from animals fed with a SC diet appeared to be unsynchronized and undirected (Fig. 2F). Conversely, RNA velocity in ARC astrocytes after 5d HFHS diet showed multiple clear directed and synchronized flows, with two major areas of origin in sub-clusters *0* and *1* and two terminal areas in sub-clusters *1* and *2* (Fig. 2F). In accordance with the transcriptional activity measured by the number of DEGs (Fig. 2D and 2E), we found less synchronized RNA velocity in cells derived from 15 HFHS diet (Fig. 2F). Interestingly, we observed that the relative number of cells in sub-cluster *1* doubled from 5d (15.2%) to 15d (31.3%) HFHS diet, whereas the cells in sub-cluster *0* nearly halved (from 22.7% to 12.2%) (Fig. 2F). This putatively resulted from the transcriptional change of cellular identity indicated by the RNA velocity after 5d of a HFHS diet, which seems to differentially affect individual astrocyte subpopulations (Fig. 2F). Next, we identified the potential driver genes of the likelihood-dependent transcriptional dynamics, and ordered them by the inferred latent time (Bergen et al., 2020) for each cluster and diet group (Fig. S2A and Table S4). Among the list of potential driver genes, we identified critical genes for astrocyte function and control of metabolism, and estimated their active expression at specific time points (Fig. S2A-B). Overall, among the drivers for transcriptional dynamics in ARC astrocytes, we found genes involved in glucose metabolism (*Aldoc*) and hormonal signaling regulation (*Pcsk1n*) in cells clustered to sub-cluster *a* in SC diet. In both HFHS diet groups, ARC astrocytes expressed specific driver genes involved in lipid metabolism (*ApoE* and *Clu*), astrocyte reactivity (*Gfap*), and mitochondrial dynamics (*Ucp2*) in cells clustered to sub-clusters *0* and *1*, which corresponded to the areas of highest RNA velocity at 5d HFHS diet (Fig. 2F and S2A-B). Together, this dataset indicates that a hypercaloric diet affects the transcriptional activity of ARC astrocytes in a time-dependent manner, with a 5 days exposure to a HFHS diet inducing a strong, yet transient, transcriptional activation.

**Figure 2.**
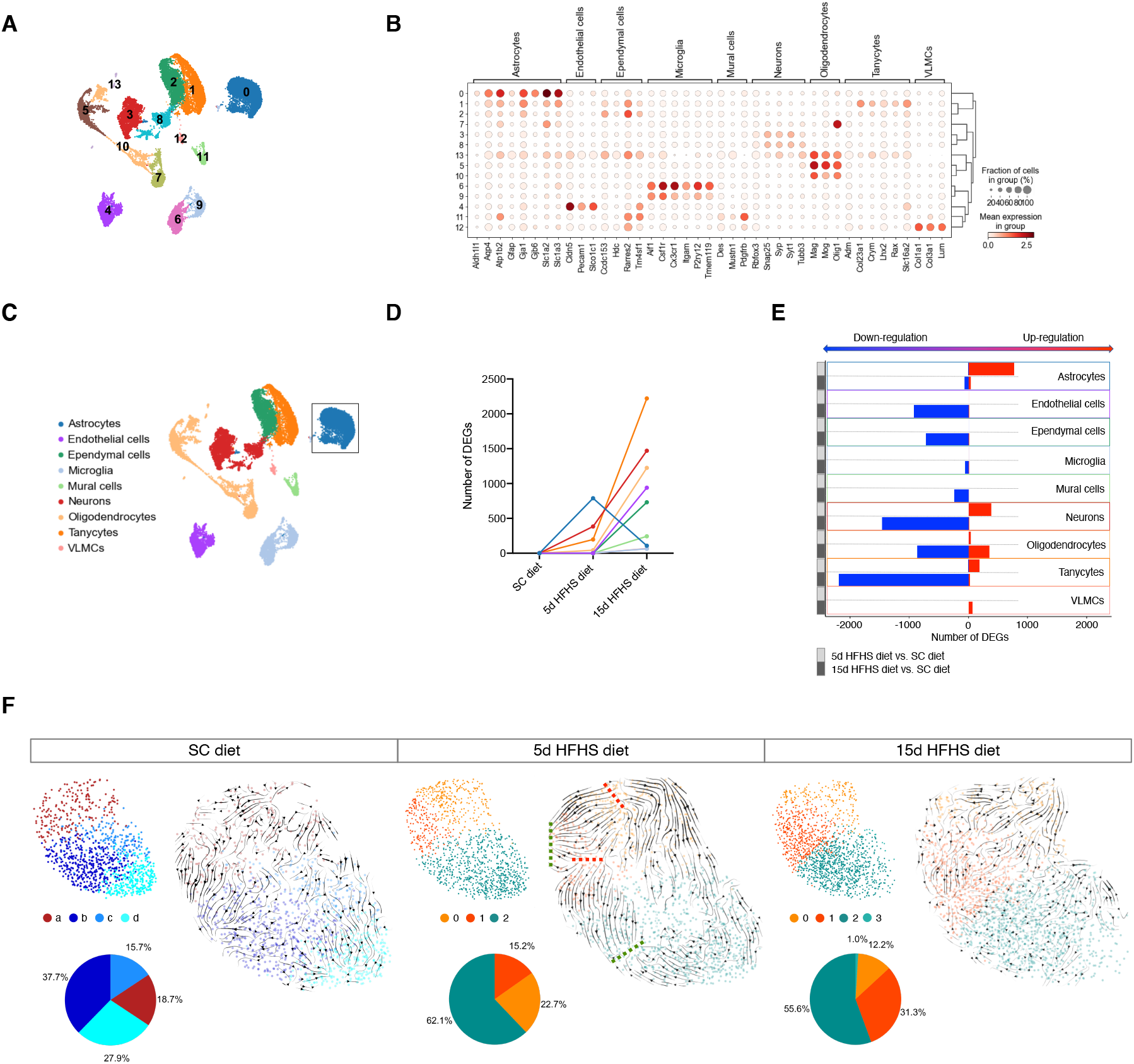
HFHS diet induces rapid transcriptional changes in astrocytes located in the ARC. **(A)** UMAP computed on full processed scRNA-Seq dataset (n = 19995 resulting after the initial filtering from 21143 cells) with annotated Leiden clusters superimposed. A median of 1802 unique transcripts was detected per each cell, resulting from the dissection of the ARC of 18 mice (6 per each diet group: SC diet, 5d HFHS diet or 15d HFHS diet). Number of cells analyzed per each diet group: SC diet = 6741; 5d HFHS diet = 5886; 15d HFHS diet = 7368. **(B)** The dot plot shows 14 clusters of interest plotted against CNS cell type-specific molecular markers. Each dot indicates the mean expression level in a particular cluster, while the size of the dots depicts the proportion of cells expressing the gene under analysis. **(C)** The color code in the UMAP embeddings indicates the cell type annotation of the clusters corresponding to **(B)**. The black square highlights the cluster identified as astrocyte-specific and used for further analyses. **(D)** The graph indicates the number of DEGs (adjusted p-value < 0.05) identified comparing the 5d or 15d HFHS diet with SC diet group per each cell type. **(E)** The numbers of DEGs shown in **(D)** are distinguished between up- (red) and down-regulated (blue) for each cell type. Light grey squares indicate DEGs identified comparing between 5d HFHS diet and SC diet groups, while dark grey squares show DEGs between 15d HFHS diet and SC diet groups. **(F)** UMAP plots of astrocytes with Leiden clustering and RNA velocity superimposed for each diet. Pie charts indicate the percentage of cells in each cluster. In 5d HFHS diet group, the red dashed lines indicate the putative starting points of RNA velocity flows, while the green dashed lines represent the terminal points. The number of cells per each sub-cluster are in SC diet: *a* = 210; *b* = 424; *c* = 176; *d* = 314; 5d HFHS diet: *0* = 282; *1* = 189; *2* = 772; 15d HFHS diet: *0* = 189; *1* = 468; *2* = 864; *3* = 15. 5d: 5 days; 15d: 15 days; Aldh1L1: Aldehyde Dehydrogenase 1 Family Member L1; DEGs: differentially expressed genes; GFAP: Glial Fibrillary Acidic Protein; HFHS: high-fat high-sugar; SC: standard chow; VLMCs: vascular leptomeningeal cells. P values for differential expression were analyzed by two-tailed Student’s *t*-test.

### HFHS diet increases the number of Aldh1L1- and GFAP-expressing astrocytes in the ARC depending on the time of diet exposure

To further study whether a calorie-dense diet changes the expression of specific molecular markers in ARC astrocytes, we examined astrocyte markers at single cell resolution in response to a HFHS diet. In SC-fed mice, the analyzed astrocyte canonical markers appeared to be expressed sparsely among astrocytes without the distinction of specific sub-clusters, suggesting that the presence of specific astrocyte-enriched molecular markers did not define separate subpopulations (Fig. S3A). Nevertheless, when we explored the temporal HFHS diet-induced changes in astrocyte specific markers (Fig. 3A and S3A), we found that astrocytes expressing Aldh1L1 and GFAP displayed a higher increase in their number after 5d of exposure to a HFHS diet versus SC diet (36.3% and 19.3% respectively), when compared to other selected markers (Fig. 3A). Conversely, other astrocyte markers analyzed (*Aqp4, Atp1b2, Gja1, Gja6, Slc1a2*, and *Slc1a3*) showed a low increase in response to 5d HFHS diet, ranged between 10% and 15%, and a moderate increase at 15d HFHS diet (25-30%) (Fig. 3A). To confirm such specific astrocytic molecular responses, we performed quantitative analysis of Aldh1L1- and GFAP-expressing populations using Aldh1L1-CreER^T2^ mice (Winchenbach et al., 2016), that were crossed to Sun1-sfGFP tagged mice, in combination with a regular GFAP antibody staining. The resulting Aldh1L1-CreER^T2^::Sun1-sfGFP mouse model allowed us to specifically visualize the green fluorescent protein (GFP) expression within the nucleus of Aldh1L1^+^ cells, circumventing the unavailability of a good anti-Aldh1L1 antibody for histological analysis. After exposure to the different diets, the tamoxifen-inducible Cre-dependent expression of GFP was induced to specifically track and visualize the diet-induced Aldh1L1 expression (Fig. S3B). Next, we proceeded to the molecular identification of these astrocytic populations in the ARC based on the single- or co-expression of GFP and/or GFAP signals. Consistently with previous observations, we found that 5d HFHS diet-fed mice exhibited a higher number of three distinct astrocyte populations when compared to SC diet-fed mice (Fig. 3B-C). Specifically, we identified astrocytes expressing: (1) Aldh1L1^+^/GFAP^-^ (Fig. 3D: white arrow); (2) Aldh1L1^-^/GFAP^+^ (Fig. 3D: white chevron); or (3) both Aldh1L1^+^/GFAP^+^ (Fig. 3D: white pentagon). Moreover, the number of Aldh1L1^+^/GFAP^-^ and Aldh1L1^+^/GFAP^+^, but not Aldh1L1^-^/GFAP^+^ cells increased in mice fed with HFHS diet for 15d, in comparison to the SC diet control group (Fig. 3C). However, the number of Aldh1L1^+^/GFAP^-^ cells in 15d HFHS diet-fed mice was significantly lower than in 5d HFHS diet-fed mice (Fig. 3C). Regarding the increase in the number of Aldh1L1^+^ cells, similar changes were also found in 5d HFHS diet-fed compared to 15d HFHS diet-fed wildtype mice using RNAscope analysis (Fig. S3C-D). Unlike Aldh1L1^+^ cells, the HFHS diet-induced increase in GFAP^+^ cells was only detected in the ARC, since a significant decrease of Aldh1L1^-^/GFAP^+^ cells was observed after HFHS diet feeding for 15d in the somatory-sensory cortex (between layers 4 and 5), and no change in the number of Aldh1L1^-^/GFAP^+^ cells was found in the hippocampal CA1 area (including stratum radiatum, stratum lacunosum-moleculare, and dentate gyrus) (Fig. S3E-H). Overall, these results indicate that a hypercaloric diet rapidly induces an increase in the number of Aldh1L1^+^ astrocytes in the different brain regions that we studied, while an increase in GFAP^+^ astrocytes was only observed in the ARC. In general, these findings reveal region-specific molecular responses of astrocytes over a HFHS diet exposure, associated with the expression of GFAP and/or Aldh1L1.

**Figure 3.**
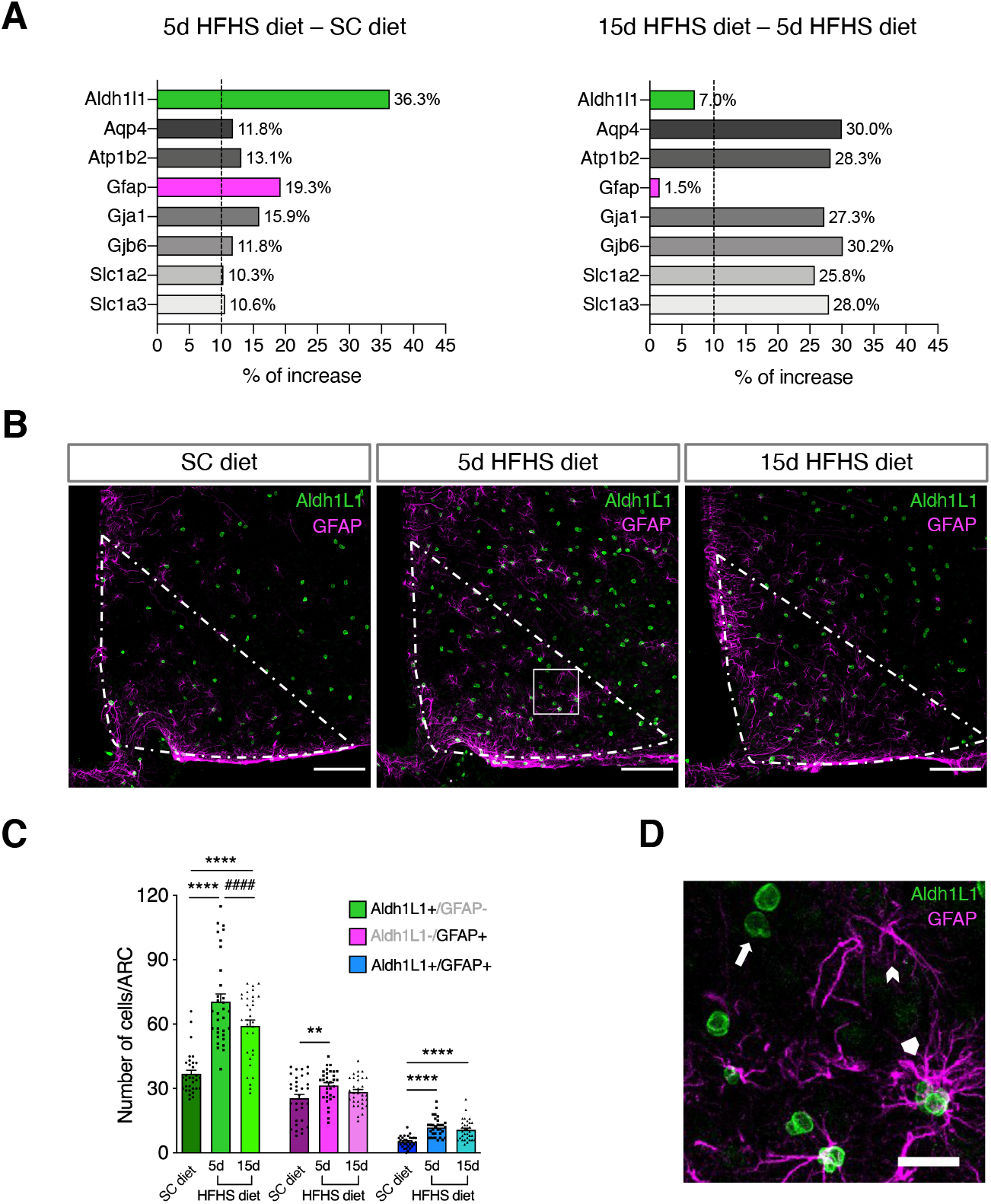
Astrocytes expressing GFAP and/or Aldh1L1 increase in number in response to a HFHS diet. **(A)** Bar graphs showing the percentage of increase of cell numbers expressing astrocyte-specific molecular markers after 5d HFHS diet compared to SC diet group and after 15d HFHS diet compared to 5d HFHS diet group, data derived from scRNA-Seq analysis. The dashed line indicates the approximative minor increase in cell number percentage (10%) when considering the molecular markers overall within the responders to 5d HFHS diet. The green (Aldh1L1) and magenta (GFAP) colored bars show the molecular markers mostly impacted by 5d HFHS diet and the least after 15d HFHS diet. **(B)** Representative images of Aldh1L1-expressing cell nuclei (green) and GFAP immunolabeling (magenta) in the ARC (area defined by segmented white line) of mice exposed to a SC diet or a HFHS diet for 5 and 15 days. White square indicates a detail of the ARC zoomed out in panel **(D)**. Scale bar = 100 µm. **(C)** Bar graph shows the number of Aldh1L1^+^/GFAP^-^ (green), Aldh1L1^-^/GFAP^+^ (magenta) and Aldh1L1^+^/GFAP^+^ (blue) cells per ARC (area of approximately 105350 μm^2^) in each experimental condition. N= 32 ARCs (derived from 4 animals, 8 ARCs per animal) analyzed per group. **p = 0.0018; ****p and ####p < 0.0001. **(D)** Zoom in the representative image from panel **(B)**. White arrow indicates an example of an Aldh1L1^+^/GFAP^-^ cell, white chevron an Aldh1L1^-^/GFAP^+^ cell, and white pentagon an Aldh1L1^+^/GFAP^+^ cell. Scale bar = 20 µm. 5d: 5 days; 15d: 15 days; Aldh1L1: Aldehyde Dehydrogenase 1 Family Member L1; ARC: arcuate nucleus of the hypothalamus; GFAP: glial fibrillary acidic protein; HFHS: high-fat high-sugar; SC: standard chow. P values for unpaired comparisons were analyzed by generalized linear model. Results are expressed as mean ± the SEM.

### HFHS diet dynamically remodels the topographical distribution of Aldh1L1- and GFAP-expressing astrocytes in the ARC

So far, our findings showed that ARC astrocytes exhibit dynamic transcriptional responses to a HFHS diet, and these changes are associated with the expression of astrocyte specific molecular markers, such as GFAP and Aldh1L1. To further investigate whether such changes occur extensively throughout the ARC or in restricted areas, we performed a topographical quantification of GFP- and/or GFAP-immunoreactive cells in the ARC of Aldh1L1-CreER^T2^:: Sun1-sfGFP mice. To this end, we defined a region of interest (ROI) corresponding to the medial ARC, with the x axis delineating the lower boundary of the ARC, the y axis delimiting the border of the third ventricle (in μm; Fig. 4A), and the origin of axes (0 μm) corresponding to the median eminence (ME); then, we measured the local density pattern of Aldh1L1^+^/GFAP^-^, Aldh1L1^-^/GFAP^+^, and Aldh1L1^+^/GFAP^+^ astrocytes (Fig. 4B). In the SC diet-fed mice, we observed that these three populations were localized in specific, highly dense areas, as depicted in Fig. 4B. Aldh1L1^+^/GFAP^-^ astrocytes showed the highest density in a single area 120 μm in the x direction from the ME, whereas Aldh1L1^-^/GFAP^+^ astrocytes were observed to be mainly concentrated in two more distal areas: one about 250 μm in x direction from the ME and a smaller one about 400 μm distal from the ME in y direction. In contrast, Aldh1L1^+^/GFAP^+^ astrocytes appeared in several smaller peaks no more than 200 μm from the ME, where the main peak almost matched the highest density area for Aldh1L1^+^/GFAP^-^ astrocytes (Fig. 4B). We next examined the effect of a HFHS diet on the topographical distribution pattern of the three identified astrocyte subtypes previously described in SC diet-fed mice. Employing a generalized linear model, we analyzed the distribution of astrocyte densities within the ARC, by dividing the total area into equal squares and then testing for cell number differences between SC and HFHS diets, both per square and overall ARC (see also Material and Methods). As before, we found an increase in the number of astrocytes in all three populations following a HFHS diet, with a prominent effect within the first 5d of hypercaloric diet exposure (Fig. 4C and S4). The increase in Aldh1L1^+^/GFAP^-^ and Aldh1L1^+^/GFAP^+^ cell numbers was more equally distributed across the ARC, whereas Aldh1L1^-^/GFAP^+^ astrocytes mainly raised in areas close to the ME (Fig. 4C and S4). Lastly, we assessed whether these astrocyte populations are spatially organized and tend to form local identical clusters in response to a HFHS diet over time. To do that, we measured the degree of spatial coherence (depicting the level of similarity between neighbors) of each astrocytic subtype in different conditions (SC diet, 5d, or 15d HFHS diet) by applying Moran I spatial autocorrelation coefficient, previously described as an indicator of the level of spatial dispersion (Schmal et al., 2017). As shown in Figure 4D, the coefficient decreased upon a HFHS diet exposure at 5 and 15 days compared to a SC diet, indicating a rising in the spatial dispersion of cellular expression over time. We also observed an increase in the occupancy of Aldh1L1^+^/GFAP^-^ specific domains in comparison to the other two populations induced by a HFHS diet (Fig. 4D). Of note, the total number of Aldh1L1^+^/GFAP^+^ astrocytes in the ARC over all diets was lower compared to Aldh1L1^+^/GFAP^-^ and Aldh1L1^-^/GFAP^+^ cells, occupying less than 1% of the total ARC area (Fig. 3C and 4D). All together, these results indicate that GFAP- and/or Aldh1L1-expressing astrocytes are organized in local clusters in the ARC, and that HFHS diet promotes the spatial dispersion of these astrocytes in a time-dependent manner.

**Figure 4.**
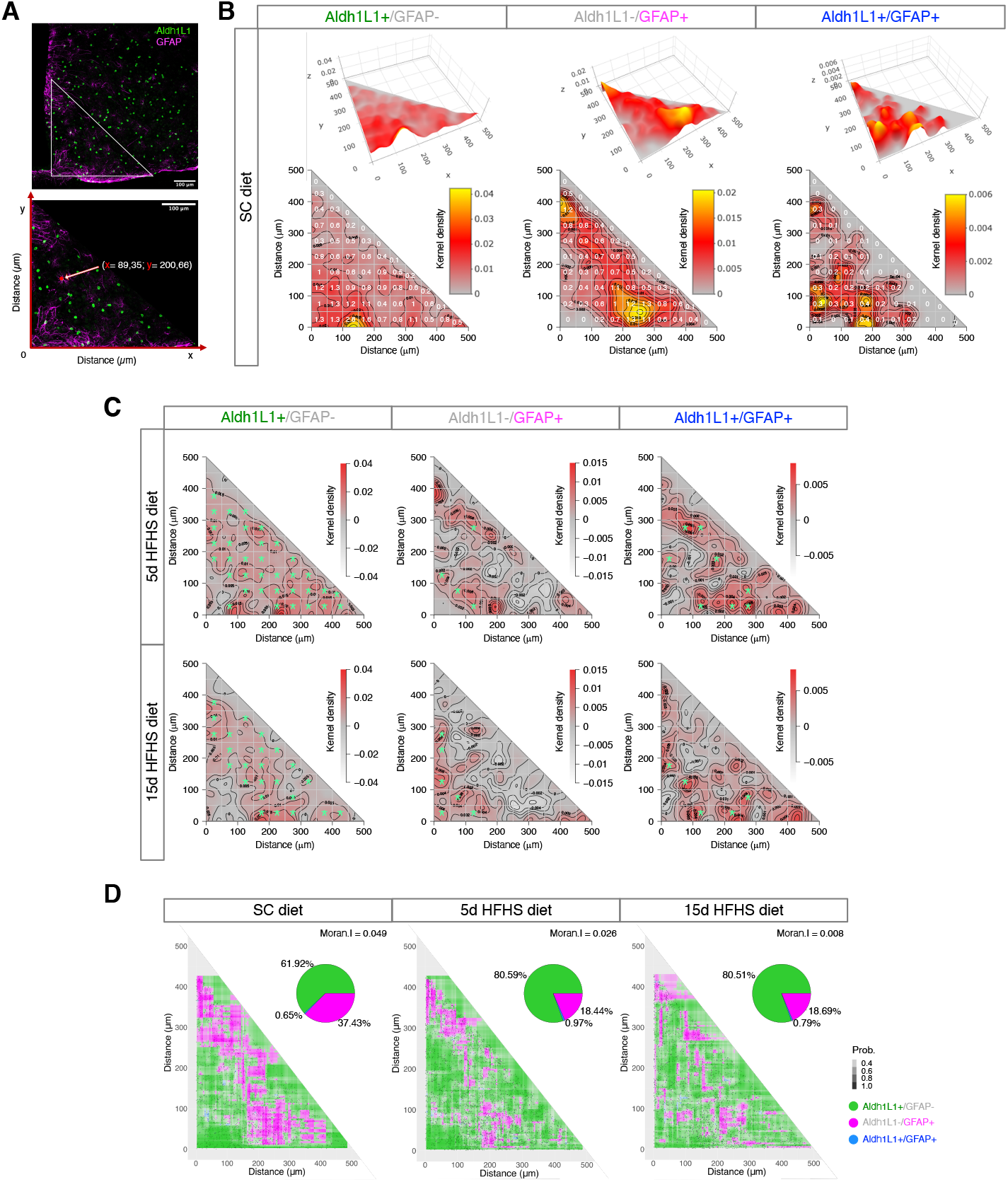
HFHS diet induces changes in the anatomical distribution of astrocytes expressing GFAP and/or Aldh1L1 in the ARC. **(A)** Representative image of Aldh1L1-expressing cell nuclei (green) and GFAP immunolabeling (magenta) enclosed in the ARC (triangle delineated with a white line; area of approximately 105350 μm^2^). The bottom picture shows an enlargement of the area adjacent to x/y axes (red), and an example of coordinates defined for one specific example cell (white arrow and red circle). Scale bars = 100 µm. **(B)** Colormaps illustrate spatial point patterns kernel density of Aldh1L1^+^/GFAP^-^, Aldh1L1^-^/GFAP^+^, and Aldh1L1^+^/GFAP^+^ astrocytes in the ARC of SC diet fed mice in 3D (upper part) and 2D (lower part). Kernel density is color coded, while contours and a white grid constituted by 50 µm^2^ small areas are shown in the 2D maps. The numbers in 50 µm^2^ squares denote the average of astrocytes over the ARCs (n = 32 from 4 animals, 8 ARCs per animal). **(C)** For each group of astrocytes expressing Aldh1L1 and/or GFAP, kernel density of patterns at SC diet group is subtracted from both 5 or 15 days HFHS diet groups. For each 50 µm^2^ area, the distribution of identified astrocytes over all the examined ARCs are compared between the SC and HFHS diet groups using the generalized linear model. The squares presenting significantly different results (p < 0.05) are marked with a green star. **(D)** Random forest classifier determines the partitioning of the feature space shared by Aldh1L1^+^/GFAP^-^, Aldh1L1^-^/GFAP^+^ and Aldh1L1^+^/GFAP^+^ astrocytes in each experimental group. Pie charts depict the percentage of coverage of each astrocytic subpopulation occupying the total area. Moran’s I spatial auto-correlation shows the correlation between Aldh1L1- and/or GFAP-expressing cells located nearby in space. SC diet: Moran I = 0.049 (p-value <0.05); 5d HFHS diet: Moran I = 0.026 (p-value <0.05); 15d HFHS diet: Moran I = 0.008 (p-value <0.05). 5d: 5 days; 15d: 15 days; Aldh1L1: Aldehyde Dehydrogenase 1 Family Member L1; ARC: arcuate nucleus of the hypothalamus; GFAP: glial fibrillary acidic protein; HFHS: high-fat high-sugar; SC: standard chow. P values for unpaired comparisons were analyzed by Friedman test.

## DISCUSSION

To date, several studies have revealed that astrocytes regionally encode their functions in the control of metabolism, depending on the specific circuits they are integrated in (Chen et al., 2016, Garcia-Caceres et al., 2016, Varela et al., 2021, Bouyakdan et al., 2019). Therefore, the study of astrocytes responses to a hypercaloric diet should be addressed preserving the precise anatomical location of these glial cells. Hypothalamic astrocytes, in particular those located in the ARC, have emerged as a particularly sensitive to diet-induced obesity (Horvath et al., 2010, Thaler et al., 2012). However, little is known about the molecular and spatial distinctions of these glial cells, which might help to determine their high sensitivity to respond to obesogenic diets. Here we reported that a high-energy food consumption triggers intra- and inter-regional molecular distinctions of hypothalamic astrocytes in mice, in contrast to other astrocytic populations and cell types in the brain. By combining bulk transcriptional and proteomic analyses, we revealed that long-term exposure to a HFHS diet induces diverse astrocyte responses, which are strongly encoded by their anatomical location. These findings indicate that astrocytes respond to a hypercaloric diet in a region-specific molecular manner, which could result from their interactions with specific local circuits (Chai et al., 2017, Morel et al., 2017, Diaz-Castro et al., 2021). Hypothalamic astrocytes, in contrast to astrocytes from other brain regions, exhibited a prominent HFHS diet-induced transcriptional regulatory switch in pathways related to hormonal and nutrient signaling processes, which have been demonstrated as critical for the regulatory role of these glial cells in mechanisms underlying nutrient and hormonal sensing in the brain (Garcia-Caceres et al., 2016, Gao et al., 2017, González-García and García-Cáceres, 2021). Given that adjacent astrocytes in the same brain region show distinct expression of molecular markers and exhibit specific capabilities to regulate the activity of neighboring neurons (Perea et al., 2014, Martín et al., 2015, Ben Haim and Rowitch, 2017), we analyzed the transcriptome of individual astrocytes to investigate their spatial and molecular diversity within the ARC. By sequencing the RNA of diverse cell populations endowed by the ARC at single-cell resolution, we found that astrocytes show the greatest gene expression changes in response to 5d HFHS diet in comparison to other cell types, including microglia and neurons. Despite the fact that our results showed minor changes in microglia of HFHS diet-fed mice, other studies have reported that a HFHS diet feeding induces a rapid reactive response in both astrocytes and microglia (Thaler et al., 2012), the latter playing a critical role in promoting the synthesis and release of inflammatory markers, which affect the functionality/activity of neighboring cells (Valdearcos et al., 2014). Given that a HFHS diet triggers fluctuations in glial reactivity associated with cellular arrangements in melanocortin neurons in the ARC (Horvath et al., 2010, Thaler et al., 2012), the lack of response to a hypercaloric diet observed in microglia could be explained by the fact that the major changes in microglia gene expression could occur at different time points than studied here. In our studies, ARC astrocytes were the major cell type affected by 5d of exposure to a HFHS diet, with a remarkable activation of their transcriptional activity that was reversed to a control-like pattern at 15d of exposure to a HFHS diet. We also observed that a HFHS diet increased the expression of two astrocyte-specific molecular markers: GFAP, which corroborates other findings from the literature (Horvath et al., 2010, Thaler et al., 2012) and Aldh1L1, that is first reported here. Interestingly, we reported that astrocytes expressing GFAP and/or Aldh1L1 populate distinct areas in the ARC, suggesting specific intra-regional functions. Moreover, a hypercaloric diet induced the increase of the area occupied by Aldh1L1-expressing astrocytes, which shows that this population responds in a different pattern than astrocytes expressing GFAP. However, it is still unknown whether the expression of these astrocytic markers is associated with functional changes in glial cells, which might in turn affect neuronal excitability and neurotransmission. The dynamic changes in the molecular profile of individual astrocytes in a spatial-dependent manner could indicate region- and network-specific astrocyte features for controlling neuronal activity according to different neuronal subtypes. Rather than acting as markers of different astrocyte subtypes, our results indicate that the expression of Aldh1L1 and/or GFAP in the ARC could entail transient states required for tissue homeostasis, without changing the common basic cellular characteristics or distinguishing between astrocytic subtypes. Whether the population of a definite area in the ARC by astrocytes expressing specific molecular markers might correlate with an adaptive synaptic rearrangement or a pathological impairment in the neurotransmitter/ion homeostasis of surrounding neurons need to be further elucidated. Hypercaloric diet primarily promotes an aberrant neuronal activity specifically in the entangled circuits of the ARC (Kälin et al., 2015, Jais et al., 2020, Jais and Brüning, 2021). Thus, the increase in GFAP- and Aldh1L1-expressing astrocytes in this area may result from (or cause) specific adaptive responses of these glial cells to alterations in different neuronal and synaptic subtypes. Further loss- or gain- of function experiments must be performed to address the question of whether the HFHS diet-induced molecular remodeling of ARC astrocytes is determinant for impairing ARC neuronal circuits sensitivity and changing feeding behavior in response to a hypercaloric diet.

### Limitations of the study

Although our study contributes to the understanding of astrocytes’ responses to hypercaloric diet, it is important to mention that the employed techniques have several limitations. In particular, ACSA2-based MACS might be biased by contamination of non-astrocyte cells. Cellular isolation preceding the RNA sequencing (scRNA & bulk RNA) potentially affects transcriptional profiles. Unlike scRNA-Seq, bulk RNA-Seq and proteomics mask cellular heterogeneity, not allowing to differentiate between specific sub-populations. In addition, our transcriptomic and proteomic measures were based on different biological samples. Nevertheless, scRNA-Seq comes with several technical limitations: low RNA capture rate is still poor compared to bulk RNA-Seq, resulting in a lower signal to noise ratios and causing very sparse expression with dropout, which might influence the interpretation of results, such as the limited starting material per cell (Kolodziejczyk et al., 2015, Potter, 2018, Chen et al., 2019). The scarcity of initial material obtained from one single animal for bulk RNA-Seq, scRNA-Seq and proteomics analyses made it necessary to pool tissues from different individuals together, which is a commonly used practice (Campbell et al., 2017, Lattke et al., 2021). In addition, by performing bulk and scRNA-seq, we lost the spatial information that might influence cellular gene expression. However, our spatial point pattern analysis comprises some limitations as well. In particular, the quantification and position determination of cells was performed on the maximum projection of 30 µm thick sections of brain tissue, which formed individual 2D plans. Additionally, projecting 3D slices to a 2D mapping might not completely represent reality, as the coordinates of cells in the z axis were not determined. Moreover, the application of this method does not exclude the possibility that cells between two different sections were not visualized and considered for the analysis. The limitations described here might be partially overcome by integrating spatial transcriptomics and scRNA-Seq techniques (Longo et al., 2021), which would provide the overall transcriptional information of each cell in a specific region, without disturbing their spatial location.

## Acknowledgements

We thank Dr. Thomas Walzthoeni for bioinformatics support provided at the Bioinformatics Core Facility (Institute of Computational Biology, Helmholtz Zentrum München), Dr. Maria Caterina De Rosa and M.D. Claudia A. Doege (Columbia University Medical Center) for providing the protocol to isolate single cells from the ARC, and Cassie Holleman, Clarita Layritz, Elisavet Lola, Nicole Klas, and Ines Kunze for their excellent technical assistance. The research leading to these results has received funding from European Research Council ERC (CGC: STG grant AstroNeuroCrosstalk # 757393), from the German Research Foundation DFG under Germany’s Excellence Strategy within the framework of the Munich Cluster for Systems Neurology (EXC 2145 SyNergy – ID 390857198), Helmholtz Excellence Network and the Deutsche Forschungsgemeinschaft (GS: SPP1757 - SA2114/2-2), and Helmholtz Association - Initiative and Networking Fund. IG-G is a recipient of a fellowship from European Union’s Horizon 2020 research and innovation program under the Marie Sklodowska-Curie actions (842080-H2020-MSCA-IF-2018). The funders had no role in study design, data collection, analyses, decision to publish, or preparation of the manuscript.

## Author contributions

LML and CG-C conceptualized all studies and designed all experiments. LML, IG-G, and BL (single-cells suspension preparation, immunohistochemistry), LML (immunohistochemistry imaging and quantification), LML and OLT (RNAscope imaging and quantification), TG, and BL (brain dissociation and MACS), MS (10x Genomics), and NK (mass spectrometry and proteomics) conducted the experiments and collected the data. GS provided Aldh1L1-CreER^T2^ mice. LML and CDBM (immunostained and fluorescent *in situ* hybridized sections) and VM (transcriptomics, proteomics, sc-RNA Seq data, spatial point pattern analysis) analyzed the data under the guidance of DL and CG-C. LML prepared and organized the figures. LML, VM, DL, and CG-C wrote the manuscript in discussion with MHT, HL, TM, and SU. All authors approved the submission.

## Supplemental Figures

**Figure S1.**
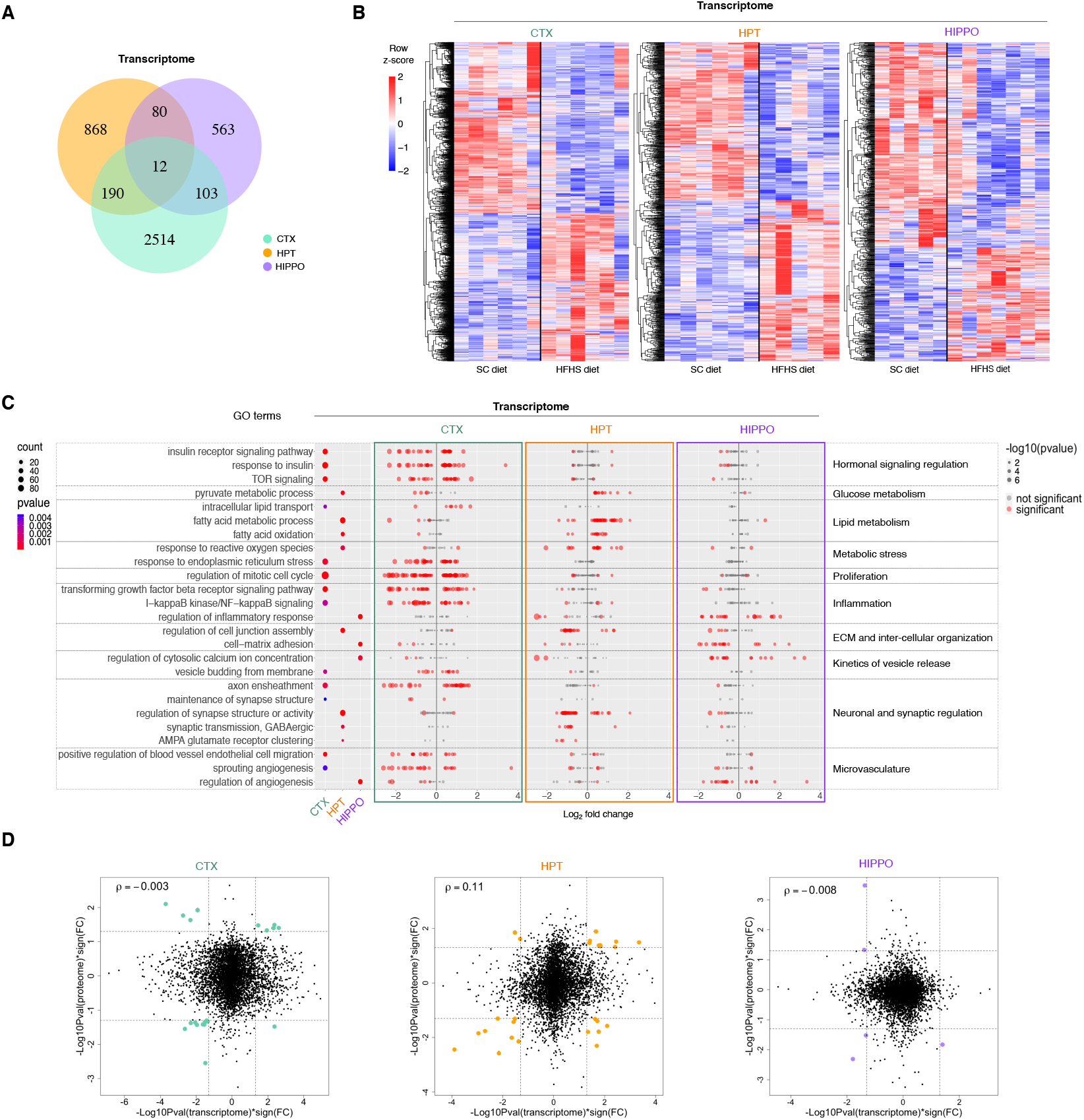
MACS-isolated astrocytes from distinct brain regions reveal transcriptional heterogeneity in response to a HFHS diet. **(A)** Venn diagram showing numbers of commonly and differentially regulated genes with p < 0.05 in ACSA2^+^ cells, comparing SC and HFHS diet groups in each brain region. **(B)** Heatmap of normalized counts intensity values obtained by unsupervised clustering of differentially expressed genes (p < 0.05) for each brain region. The color code in the heatmap indicates row z-score normalized expression values. Samples derived from SC or HFHS diets-fed animals are separated by a black line. Each column represents one sample, and each sample is formed by ACSA2^+^ cells derived from 2 animals. **(C)** The dot plot illustrates manually selected GO terms of interest enriched for DEGs identified in brain regions comparing SC and HFHS diet groups, in transcriptomics (p < 0.05). The left panel shows the dot plot of the selected GO enriched biological process terms for each brain tissue. The dot size indicates the number of DEGs, while the color depicts statistical significance of the enrichment. The dot plots in the three adjacent panels correspond to the three different brain regions analyzed, where each gene mapped to a specific pathway is represented by a single dot. The dot color refers to the presence (red) or absence (grey) of a statistically significant value, and the dot size indicates the level of significance. The x-axis represents log_2_ fold change. (D) For each brain region, -log_10_(p-value) multiplied with sign of the fold change of proteins are plotted against the transcriptomics. The colored dots represent the significant genes (p-value < 0.05). In the top left corner of each panel, Spearman’s rank correlation coefficient ρ is plotted. CTX: cortex; ECM: extracellular matrix; FC: fold change; GO: gene ontology; HFHS: high-fat high-sugar; HIPPO: hippocampus; HPT: hypothalamus; SC: standard chow; ρ: Spearman correlation coefficient. P values for differential gene expression analysis were analyzed by Wald test.

**Figure S2.**
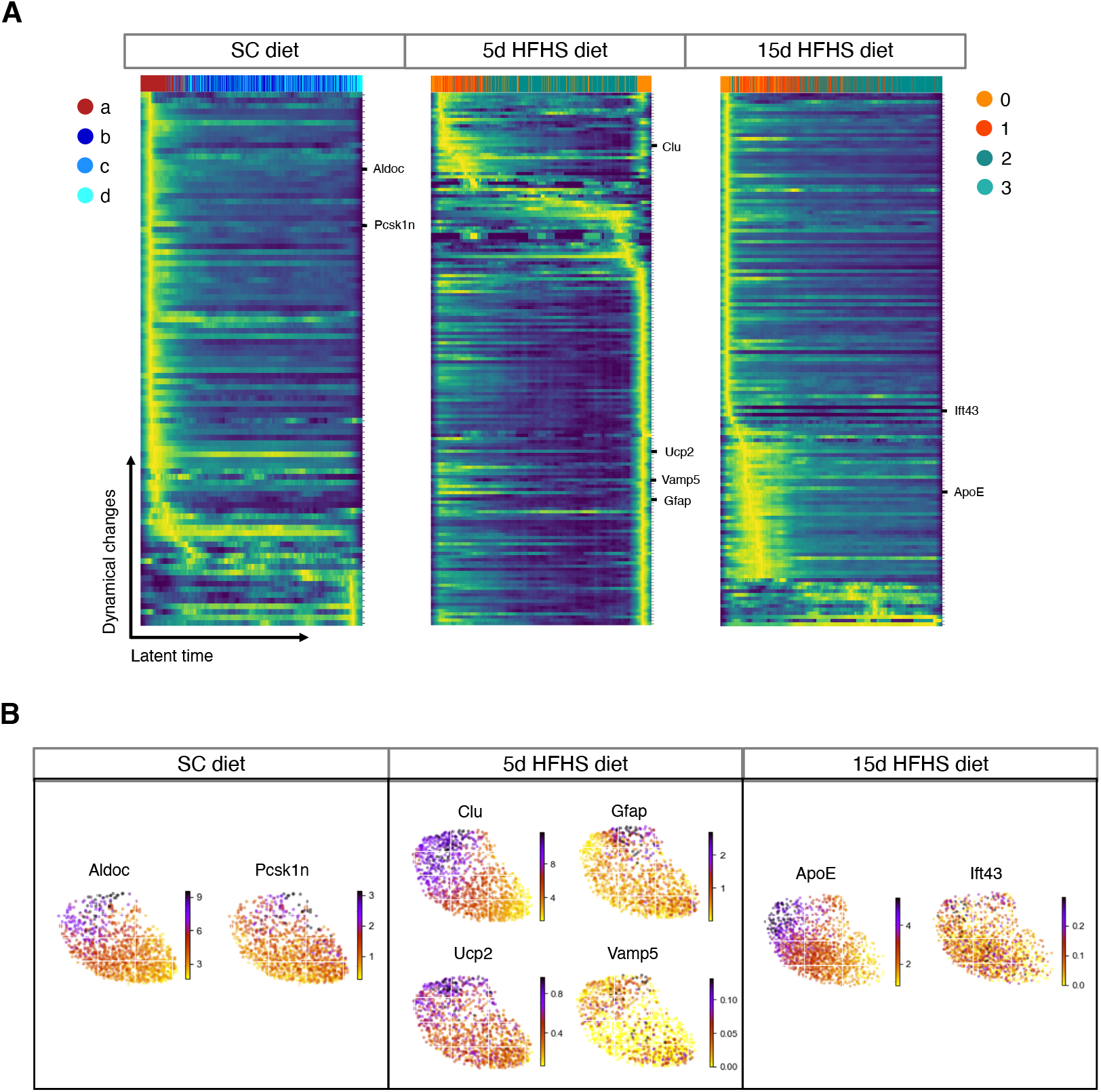
HFHS diet induces rapid transcriptional changes in astrocytes located in the medial ARC. **(A)** Heatmaps for each diet depict the RNA velocity potential driver’s expression plotted against the latent time, while the colors on the top indicate the cluster affiliation of each cell. Some genes of interest are indicated on the right side of the plots. **(B)** UMAP plots showing the expression of the potential driver genes indicated in the heatmaps for each diet. 5d: 5 days; 15d: 15 days; HFHS: high-fat high-sugar; SC: standard chow.

**Figure S3.**
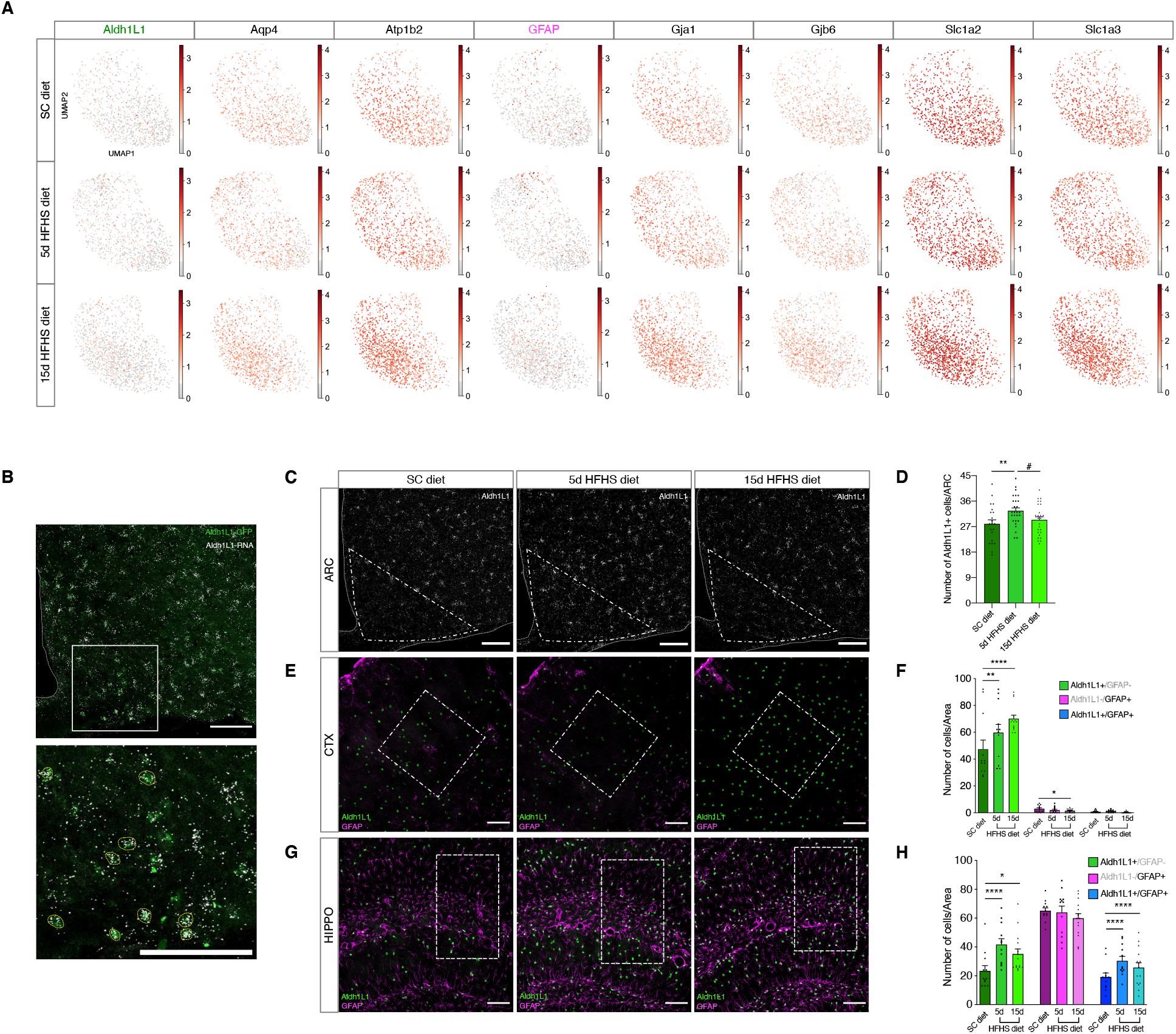
Astrocytes expressing GFAP and/or Aldh1L1 increase in their number in response to a HFHS diet. **(A)** UMAPs showing the expression of selected astrocyte canonical markers in astrocyte population for each diet condition. **(B)** Representative picture (top) and enlargement of the area enclosed in the white square (bottom) of co-localization (yellow circles) between Aldh1L1 RNA (white) and protein (green) expression. Scale bars = 100 µm. **(C)** Representative pictures of Aldh1L1-RNA (white) expression in the ARC (area of approximately 105350 μm^2^ defined by segmented line) of mice exposed to a SC or a HFHS diet for 5 and 15 days. Scale bar = 100 µm. **(D)** Bar graph shows the number of cells positive for Aldh1L1-RNA in the ARC for each experimental condition. SC diet: n = 21 (from 4 mice); 5d HFHS diet: n = 27 (from 5 mice); 15d HFHS diet: n = 26 (from 5 mice). **p = 0.0059; #p = 0.0355. **(E)** Representative images of Aldh1L1-expressing cell nuclei (green) and GFAP immunolabeling (magenta) in a defined cortical region (area of approximately 144334 μm^2^ delimited by segmented white line) of animals fed with a SC or a HFHS diet. Scale bar = 100 µm. **(F)** The bar plot shows the number of Aldh1L1^+^/GFAP^-^ (green), Aldh1L1^-^/GFAP^+^ (magenta) and Aldh1L1^+^/GFAP^+^ (blue) cells per cortical region of interest under different experimental conditions. SC diet, 5d HFHS diet: n = 12 (from 4 animals); 15d HFHS diet: n = 15 (from 5 animals). **p = 0.0019; ****p < 0.0001; *p = 0.027. **(G)** Representative images of Aldh1L1-expressing cell nuclei (green) and GFAP immunolabeling (magenta) in a defined hippocampal region (area of 96138 μm^2^ delimited by segmented white line) of animals exposed to different diets. Scale bar = 100 µm. **(H)** Bar graph indicates the number of Aldh1L1^+^/GFAP^-^ (green), Aldh1L1^-^/GFAP^+^ (magenta) and Aldh1L1^+^/GFAP^+^ (blue) cells per hippocampal region of interest of mice fed with a SC or a HFHS diet. SC diet, 5d HFHS diet: n = 12 (from 4 animals); 15d HFHS diet: n = 15 (from 5 animals). Aldh1L1^+^/GFAP^-^ SC diet vs. 5d HFHS diet: ****p < 0.0001; *p = 0.012. 5d: 5 days; 15d: 15 days; Aldh1L1: Aldehyde Dehydrogenase 1 Family Member L1; ARC: arcuate nucleus of the hypothalamus; GFAP: glial fibrillary acidic protein; HFHS: high-fat high-sugar; SC: standard chow; UMAP: uniform manifold approximation and projection. P values for unpaired comparisons were analyzed by generalized linear model. Results are expressed as mean ± the SEM.

**Figure S4.**
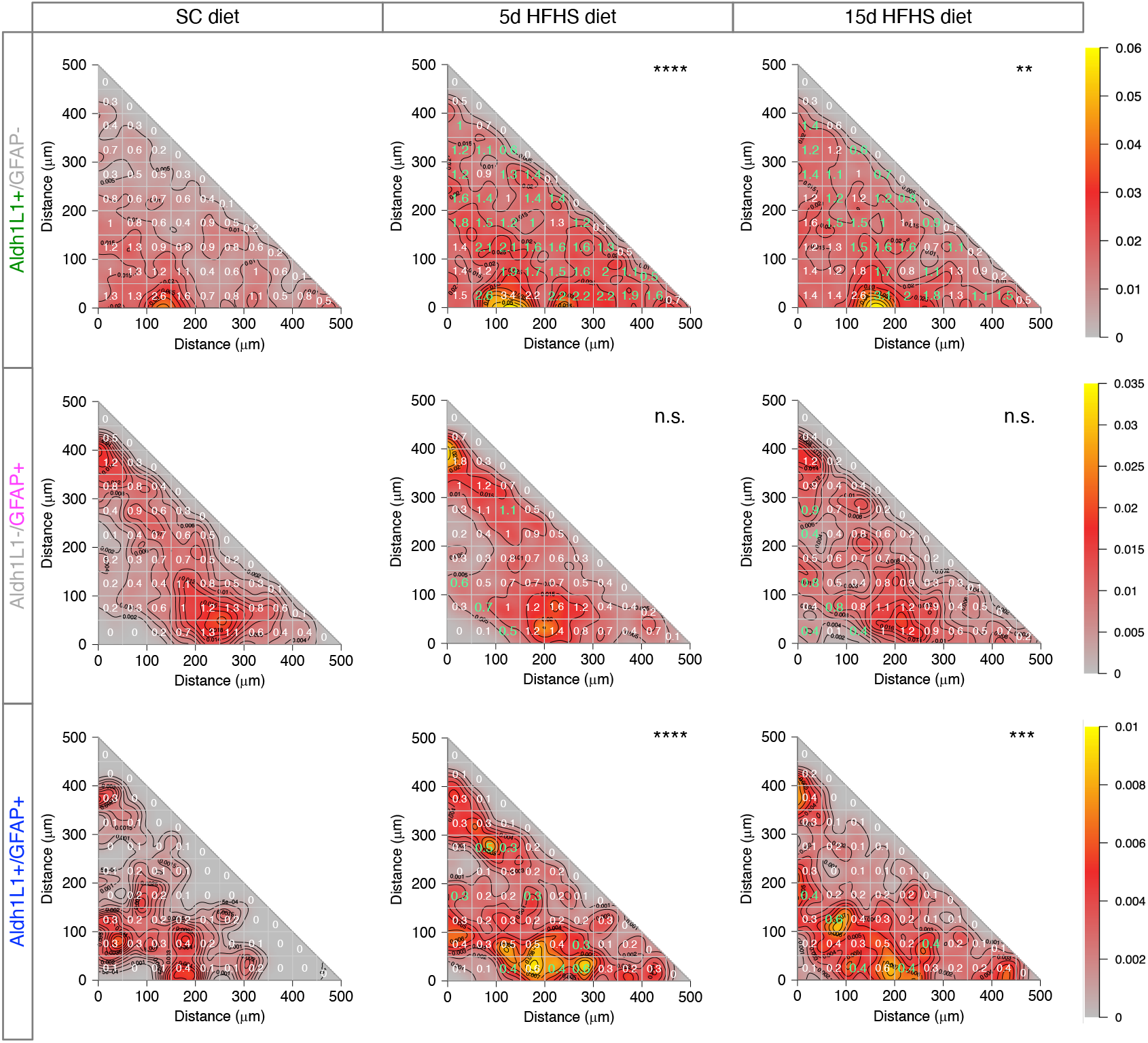
HFHS diet induces changes in the anatomical distribution of astrocytes expressing GFAP and/or Aldh1L1 in the ARC. Each plot illustrates spatial point patterns kernel density of Aldh1L1^+^/GFAP^-^, Aldh1L1^-^/GFAP^+^, and Aldh1L1^+^/GFAP^+^ astrocytes in the ARC from SC and HFHS diet-fed mice. The plots comprise kernel density color coded maps with contours and a white grid constituted by 50 µm^2^ small areas. In each square, the number indicates the average of astrocytes (n = 32 from 4 animals), and are colored in green when the comparison between SC and HFHS diet groups produce a significant value (p < 0.05). ****p < 0.0001; **p = 0.0042; ***p = 0.00053. 5d: 5 days; 15d: 15 days; Aldh1L1: Aldehyde Dehydrogenase 1 Family Member L1; GFAP: glial fibrillary acidic protein; HFHS: high-fat high-sugar; n.s.: not significant; SC: standard chow. P values for unpaired comparisons were analyzed by generalized linear model.

## Supplemental tables

**Table S1:** Lists of overlapping and non-overlapping DEGs between brain regions, and complete lists of GO pathways enriched in transcriptomics data per brain region.

**Table S2:** Complete lists of GO pathways enriched in proteomics data per brain region.

**Table S3:** Complete lists of pathways enriched both in up- or down-regulated features in transcriptomics and proteomics data per brain region.

**Table S4:** List of DEGs, divided between up- and down-regulated, in cluster 0 (astrocytes), and lists of cluster-specific potential driver genes of RNA velocity.

## STAR Methods

### Mice

All animal experiments were approved and conducted under the guidelines of the Institutional Animal Care and Use Committee of the Helmholtz Center Munich, Bavaria, Germany. Only male animals were used for all experiments, and were all housed in groups under a 12-h light/12-h dark cycle at 22 ± 2°C with free access to food and water, and maintained on a pelleted SC diet (5.8% fat; Harlan Teklad; #LM-485). Mice were randomized and evenly distributed among test groups according to age and body weight. C57BL/6JRj were obtained from Janvier Lab (Saint-Berthevin Cedex, France), and exposed to a HFHS diet (58% kcal fat w/sucrose; Research Diets; #D12331), or maintained on SC diet for a designated amount of time, accordingly to each experiment. Tamoxifen-inducible Aldh1L1-CreER^T2^::Sun1-sfGFP mice were generated by crossing Aldh1L1-CreER^T2^ (Tg(Aldh1l1-cre/ERT2)02Kan) mice (Winchenbach et al., 2016) (gently provided by G. Saher, Max Planck Institute, Göttingen), with Sun1-sfGFP (B6;129-Gt(ROSA)26Sor^tm5(CAGSun1/sfGFP)Nat^/J) mice (Mo et al., 2015) (The Jackson Laboratory, JAX 021039) to induce the green fluorescent protein (GFP) expression specifically in the nuclei of Aldh1L1-expressing cells. Tamoxifen (Sigma-Aldrich; #T5648) was injected once a day (1 mg/100 µl, *i*.*p*.) in the last two days of experimental diet exposure. Tamoxifen was dissolved in sunflower oil, then sterilized by filtration, and stored in aliquots at -20°C, protected from light, until use.

### Brain homogenization and magnetic-activated cell sorting (MACS)

C57BL/6JRj mice were exposed to a SC or a HFHS diet at 8-9 weeks of age for 4 months, and then sacrificed for tissue collection. Hypothalami, hippocampi, and half cortices derived from 2 animals (for bulk RNA-Seq) and from 4 animals (for protein mass spectrometry) were rapidly dissected and pooled together in distinct tubes prefilled with cold D-PBS. Tissue dissociation was performed following manufacturer’s instructions (Adult Brain Dissociation Kit, Miltenyi BioTec; #130-107-677). The resulting cell suspensions were incubated with magnetic microbeads coupled with an anti-ACSA2 antibody (Anti-ACSA-2 MicroBead Kit, Miltenyi BioTec; #130-097-678), and labeled cells were sorted via MACS, according to commercially available manufacturer’s protocol (Anti-ACSA-2 MicroBead Kit, Miltenyi BioTec; #130-097-678). In particular, for hypothalamic and hippocampal samples (pre-labeled number of cells < 10^7^), standard reagent volumes and MS columns were used, while for cortical samples (pre-labeled number of cells > 10^7^), reagent volumes were scaled up accordingly and LS columns were used. The resulting ACSA2^+^ and ACSA2^-^ fractions were collected in cold PBS and centrifuged at 3000 x g at 4°C for 10 min, and the pelleted cells stored at -20°C (for protein isolation) or -80°C (for RNA extraction) until use.

### RNA extraction and sequencing

The RNA extraction from ACSA2^+^ cells was performed (RNeasy Micro Kit; Qiagen; #74004) following the manufacturer’s protocol. Next, SMART-Seq® v4 Ultra® Low Input RNA Kit for Sequencing (Takara; #634891) was used to generate and amplify cDNA, according to the instructions provided with the kit. The bulk RNA-Seq was performed by Novogene (Europe, United Kingdom).

### Protein isolation and mass spectrometry (MS)

The isolation of proteins from ACSA2^+^ and ACSA2^-^ cells was performed by resuspending the pellet in 80 μl of cold RIPA buffer (Sigma-Aldrich; #R0278-500) containing a cocktail of protease and phosphatase inhibitors (Thermo Fisher Scientific Inc.; #78446) in a 1% v/v dilution, and the sample was homogenized. The cell debris were then pelleted at 13000 x g and 4°C for 10 minutes, and the protein-rich supernatant was collected. 2µg of protein was heated for 5 min at 95°C, and sonicated in 4% sodium deoxycholate (SDC), 100 mM Tris pH 8.5. After alkylation and reduction with 10 mM tris-(2-carboxyethyl)-phosphin-hydrochlorid (TCEP), 40 mM 2-chloroacetamide (CAA), the protein was digested overnight with 1:50 LysC and Trypsin at 37°C. The digested peptides were acidified and purified on SDB-RPS STAGE tips (3M Empore) (Kulak et al., 2014). Samples were concentrated in a SpeedVac for 40 min at 45°C and dissolved in 10µl MS loading buffer (2% acetonitrile (ACN), 0.1% trifluoroacetic acid (TFA)). For MS analysis, 1 µg of peptides were loaded onto a 50-cm column with a 75µM inner diameter, packed in-house with 1.9µM C18 ReproSil particles (Dr. Maisch GmbH) at 60°C. The peptides were separated by reversed-phase chromatography on a 120 min gradient (5-30% buffer B over 95 min, 30-60% buffer B over 5 min followed by washout) using a binary buffer system consisting of 0.1% formic acid (buffer A) and 80% ACN in 0.1% formic acid (buffer B) at a flowrate of 350nl on an EASY-nLC 1200 system (Thermo Fisher Scientific). MS data were acquired on a Quadrupole-Orbitrap instrument (Q Exactive HF-X, Thermo Fisher Scientific) using a data dependent top-15 method with maximum injection time of 20 ms, a scan range of 300–1650Th, and an AGC target of 3e6. Sequencing was performed via higher energy collisional dissociation fragmentation with a target value of 1e5, and a window of 1.4Th. Survey scans were acquired at a resolution of 60,000. Resolution for HCD spectra was set to 15,000 with maximum ion injection time of 28 ms and an underfill ratio of 20%. Dynamic exclusion was set to 30 s.

### Bulk transcriptomics and proteomics analyses

The raw transcriptomics sequencing reads were first evaluated using FastQC (Andrews, 2010), and after passing quality control, the expression of the transcripts was aligned and quantified against the reference index created based on the GRCm38 genome using Salmon (Patro et al., 2017), with default parameters. These transcript abundances were then imported into R (v4.0.3, R Core Team, 2019), and summarized with tixmeta (Love et al., 2020). Employing DESeq2 (Love et al., 2014), gene expression levels were normalized with median ratio method, and principal components were calculated on variance stabilizing transformed counts, while DEGs (Differentially Expressed Genes) comparing SC and HFHS diet groups for each brain region were identified using standard parameters and simple two-group comparison Wald test.

For proteome analysis, the MaxQuant software version 1.6.7.0. (Tyanova et al., 2016) was used for processing raw mass spectrometry data. Missing values were replaced from normal distribution using the data imputation feature from the Perseus software (Max Planck Institute of Biochemisty, Munich, Germany) (Tyanova et al., 2016). Significant differences in the protein levels comparing SC and HFHS diet groups for each brain region were evaluated using Student t-test.

Both for transcriptomics and proteomics data, principal component analysis plots were generated using ggplot2 (Wickham, 2016) package from R. Differential expression analysis results were illustrated with Venn diagrams using R-package VennDiagram (Chen and Boutros, 2011). The expression profile of DEGs over the samples was shown in the heatmap using R-package pheatmap (Kolde, 2015). Further, GO enrichment analyses were performed, and the results were illustrated using clusterProfiler (Yu et al., 2012) and ggplot2 (Wickham, 2016) R-packages. Integrated analysis plot of transcriptomics and proteomics data, as well as the calculation of Spearman’s rank correlation coefficient were obtained using R Base package. GO enrichment analysis for co-upregulated and co-downregulated features of each brain region was performed using clusterProfiler (Yu et al., 2012) package from R.

### Preparation of single-cells suspension from the ARC

At the weaning time, C57BL/6JRj mice were divided into three groups: the first and the second group received SC diet or HFHS diet for 15d, while the third group was fed initially with SC diet for 10 days, and successively with HFHS diet for 5d. Afterwards, all animals were sacrificed and brains rapidly extracted and positioned with the dorsal surface at the bottom of a chilled rodent brain matrix (ASI-instruments). Next, a 1 mm-thick coronal section made approximately at the middle of the hypothalamus (about 1 mm from the optic chiasm) was collected. While keeping the brain slice on ice, the ARC was isolated with a tissue puncher (1 mm diameter, #Thermo Fisher Scientific Inc.) and collected into cold DMEM/F12 medium (Thermo Fisher Scientific Inc.; #11320033). The ARC samples derived from 6 mice were pooled together and washed once with cold D-PBS containing 0.05 g/ml of Trehalose (Sigma-Aldrich; #T0167). The buffer was then discarded and the ARC samples incubated for 16 min at 37°C with 200 µl of 4 mg/ml collagenase/dispase solution (Sigma-Aldrich; #10269638001) supplemented with 0.05 g/ml of Trehalose. After the incubation period, the enzymatic solution was carefully removed and the tissue was incubated for 12 min at 37°C with 200 µl of papain/DNAse solution (prepared following steps 2 and 3 from the kit’s catalogue; Worthington Biochemical Corporation; #LK003150) supplemented with 0.05 g/ml of Trehalose. The enzymatic solution was then delicately removed and the tissue resuspended in 200 µl of D-PBS supplemented with 0.05 g/ml of Trehalose and RNAseOUT (1/1000; Thermo Fisher Scientific Inc.; #10-777-019), and gently tritured until the solution appeared milky and devoid of visible tissue pieces. The resulting cell suspension was filtered through a 40 µm cell strainer, and cells were pelleted by centrifugation for 8 min at 300 x g and 4°C in 40 ml of D-PBS containing 2% bovine serum albumin (BSA) (Miltenyi BioTec; #130-091-376). Cells were then washed one time by centrifugation (300 x g, 10 min, 4°C) in D-PBS containing 0.04% BSA, and resuspended in 40 µl of the same fresh solution. The final number and viability of the cells was assessed with a hemocytometer after staining with trypan blue. The final volume was adjusted to obtain a concentration of 1000 cells/µl, and cells were immediately processed for scRNA-Seq library preparation with a target recovery of 10000 cells. Chromium Single Cell 3□ Reagent Kits v3.1 (10x Genomics, #PN-1000121) was used to prepare libraries, according to the manufacturer’s instructions. Libraries were pooled and sequenced according to 10x Genomics’ recommendations on an Illumina NovaSeq6000 system with a target read depth of 50000 reads/cell.

### Single-cell transcriptomics analysis

The sequence and annotation files for the mouse genome GRCm38 assembly and annotation release 100 from Ensembl were used for the alignment of the reads. The Cell Ranger pipeline (version 3.1.0, from 10X Genomics, Pleasanton, CA) was run with the command ‘cellranger mkref’ to create an index of the genome. For the alignment of the reads, generation of QC metrics, estimation the numbers of valid barcodes, and generating the count matrices the Cell Ranger software were run with the command ‘cellranger count.’ The command was executed with standard parameters, while using 10000 for number of expected cells and “SC3Pv3” for chemistry parameter arguments. An anndata object was created using python package Scanpy (v1.4.4) (Wolf et al., 2018).

The single-cell transcriptomics analysis was performed using scanpy version 1.7.1 (Wolf et al., 2018). The filtering of the cells was based on a minimum UMI count of 500. Cell-wise gene expression was normalized applying the default setting to a total count of 1e4 by linear scaling. On the normalized data, we performed a PCA, while 50 principal components (PC) were used to compute a k-nearest neighbor (kNN) graph (k = 100, method = umap). The Leiden clustering (resolution = 0.3, flavor = vtraag) was computed based on the kNN graph. Further, we performed Leiden clustering and cell type identification based on canonical markers. Cluster 0 highly expressed astrocytic markers, which we considered for further investigation. Cells were processed again by filtering for a minimum UMI count of 500, then normalizing to 1e4 counts, transformation log(x+1) of the data was followed by PCA with 50 PCs. Once again, a kNN graph (k = 100, method = umap) was generated based on the PC space. UMAP plots were generated for each diet with annotated Leiden clustering superimposed, while the barplot chart (matplotlib) (Hunter, 2007) shows the absolute number of cells per cluster. Differential expression analysis was performed using Student’s t-test to compare SC diet with 5 and 15 days HFHS diet groups for each cell type, plotting the number of DEGs, as well as the number of up- and down-regulated DEGs. To further assess the underlying kinetics of gene expression of astrocytes, we performed RNA velocity estimation (La Manno et al., 2018) implemented as a stochastic version in the scVelo python package (https://github.com/theislab/scvelo). We generated a loompy file, and extracted spliced and unspliced reads using the velocyto pipeline (http://velocyto.org). With Scanpy, the file was read into an AnnData object for downstream analysis. Employing scvelo python pipeline, the calculation of RNA velocity values was performed for each gene in each cell, and the graph was then used to project the velocities into the low dimensional UMAP embedding. Using the same pipeline, the potential driving genes, which halve high likelihoods in the dynamic models and shows pronounced dynamic behavior, were identified for each diet.

### Fluorescence *in situ* hybridization

To validate the co-localization of Aldh1L1 RNA and protein, we injected one 12 week-old Aldh1L1-CreER^T2^::Sun1-sfGFP mouse with tamoxifen for two consecutive days, and sacrificed it three days later. To assess the HFHS diet-induced changes in the levels of Aldh1L1-RNA in the ARC, we provided a SC diet or a HFHS diet to 10-week old C57BL/6JRj mice for 5 or 15 days. All mice were perfused with 0.9% w/v NaCl and a cold 4% solution of PFA in 1x phosphate-buffered saline (PBS) (pH 7.4). Fluorescent *in situ* hybridization was then performed using the RNAscope® Multiplex Fluorescent Reagent Kit v2-Mm (ACD, Advanced Cell Diagnostic, #323100) and following the manufacture’s protocol for fixed-frozen sections (10 µm). Probe targeting *M. musculus* Aldh1L1 mRNA (RNAscope® Probe Mm-Aldh1l1; ACD, #405891), and Opal 570 Fluorophore Reagent (1/1000; Akoya Biosciences) were used. To retrieve the GFP signal, slides were successively incubated with primary chicken anti-GFP antibody (1/500 in 0.1% bovine serum albumin (BSA) in TBS, OriGene, #AP31791PU-N) for 2h, and then with secondary rabbit HRP-conjugated anti-chicken IgG (H&L) antibody (1/1000 in 0.1% bovine serum albumin (BSA) in TBS, Rockland Immunochemicals, #603-4302) for 30 min, and finally with Opal 520 Fluorophore Reagent (1/1000; Akoya Biosciences) for 15 min, mounted and stored at 4°C until further use. The samples were imaged using a confocal microscope (Leica TCS SP8) with a 20X magnification, as a z-stack with a 0.5 µm z-step size, applying the same acquisition parameters to all samples. Images were then analyzed on Fiji software on maximum intensity Z-stack projections. In particular, the same ROI used for IHC analysis was applied to the figures in order to define the ARC. The quantification of the number of Aldh1L1^+^ cells was done manually, as the specific location of Aldh1L1 RNA in the cytoplasm allowed to recognize the cellular shape, especially when highly dense. Areas with low and sparse RNA expression were not counted as cells, and ignored.

### Immunohistochemistry

A SC diet or a HFHS diet was provided to 10 week-old Aldh1L1-CreER^T2^::Sun1-sfGFP mice for 5 or 15 days, and tamoxifen was injected the last two days of experimental diet exposure to selectively induce the expression of GFP in the nuclei of Aldh1L1-expressing cells. Three days after the last tamoxifen injection, mice were perfused with 0.9% w/v NaCl and a cold 4% solution of paraformaldehyde (PFA) (Carl Roth) in 1x PBS (pH 7.4). Brains were post-fixed in 4% PFA solution at 4°C overnight, and then equilibrated for 24-48h with 30% sucrose in 1x tris-buffered saline (TBS) (pH 7.2) at 4°C. Next, 30 µm coronal sections were cut on a cryostat, washed and blocked for 1h with a blocking buffer containing 0.25% gelatin and 0.2% Triton X 100 dissolved in 1x TBS. Slices were then incubated overnight at 4°C with primary antibodies diluted in the blocking buffer: rabbit anti-GFAP (1/1000, Dako, #Z0334), chicken anti-GFP (1/500, OriGene, #AP31791PU-N). On the second day, sections were rinsed four times with 1x TBS, and incubated for 2h at RT with corresponding fluorescent secondary antibodies diluted in the blocking buffer: Alexa Fluor 647 donkey anti-rabbit IgG (H + L; 1/1000, Invitrogen, #A-31573) and Alexa Fluor 488 goat anti-chicken IgG (H + L; 1/1000, Invitrogen, #A-11039). After several TBS washes, slices were mounted on glass slides and stored at 4°C for at least one night before imaging.

### Imaging and quantification

All images were captured with a confocal microscope (Leica TCS SP5) at 20X magnification. The final z-stack generated was achieved at constant 2 µm step size with a total of 17-22 optical slices. The assessment of the numbers and position coordinates of Aldh1L1-expressing and GFAP-ir cells was performed on maximum intensity z-stack projections by using Fiji software. Only coronal slices located between Bregma -1.58 mm and -1.82 mm were included in the analysis. A ROI corresponding roughly to the ARC was drawn based on the Allen Mouse Brain Atlas (Lein et al., 2007, Atlas, 2011), with a total area of approximately 105350 μm^2^. The ROI corresponded to a right-angled triangle, with one approx. 490 μm long cathetus defining the lower boundary of the ARC (x axis), and the other cathetus approx. 430 μm long located at the border of the third ventricle (y axis). The same ROI was used for the analysis of all ARCs, and its positioning in the image was kept as consistent as possible between pictures. The number of cells and their position within the ROI were quantified manually and blindly. In particular, the defined x/y coordinates corresponded to the center of Aldh1L1+ nuclei, and to the converging point between main processes of GFAP-ir cells. Cortical and hippocampal analyses were performed on the same brain sections used to assess the number and position of astrocytes in the ARC. The ROIs defined for the analysis of cortical and hippocampal astrocytic numbers were squares with an area of approximately 144334 μm^2^ (cortex) and 96138 μm^2^ (hippocampus), positioned consistently throughout the slices in order to include layers 4 and 5 (cortex), and stratum radiatum, stratum lacunosum-moleculare, and dentate gyrus (hippocampus). The cell numbers assessment in all brain areas was performed consistently.

### Spatial point pattern analysis

All the cells identified over the slices were projected onto the 2D coordinates of the ROI. Coordinates for each cell were used for visualization and analysis of spatial point patterns. For the illustration, we generated kernel density based spatial point pattern 3D and 2D plots, using R-packages plotly (Sievert, 2020) and statspat (Baddeley and Turner, 2005), respectively. For overall significance testing of spatial point patterns, comparing SC diet with 5d or 15d HFHS diet, generalized linear model was employed for fitting the data *y* ∼ *β*_0_ + *β*_1_ * *diet* + *β*_2_ * *mouse*, where the mouse variable represents the random effect and diet the fixed effect, while the p-values were obtained using asymptotic Wald test, from R-package lme4. Using the same statistical test, the difference between the squares was investigated, comparing the number of cells identified on every slice between SC diet and 5d or 15d HFHS diet. For random forest classification, R-packages randomForest (Liaw and Wiener, 2002), mlr3, mlr3learners, and mlr3viz (Becker et al., 2021) were used, while for Moran I spatial autocorrelation calculation and testing ape R-package was employed (Paradis and Schliep, 2019).

### Statistical analyses

Statistical and computational analyses were performed using GraphPad Prism, R (v4.0.3, R Core Team, 2019), python (v3.9.2, http://www.python.org) and Perseus (v1.6.7.0) (Tyanova et al., 2016). For the assessment of astrocytic numbers, a generalized linear model together with asymptotic Wald test was used to compare two groups. P values < 0.05 were considered significant. All results are presented as mean ± SEMs.

